# Motivational modulation of flight speed in foraging bumblebees

**DOI:** 10.1101/2024.03.06.583792

**Authors:** Romain Willemet

## Abstract

The set of cognitive mechanisms employed by foraging bees is the subject of intense research due to the importance of pollination as an ecological process and as a study system in comparative psychology. Yet, our understanding of how reward valuation influences flight behaviour, and, more generally, the motivational control of behavioural vigour, remains limited. Here, the flight behaviour of individual bumblebees *Bombus impatiens* was recorded during a series of foraging trials in which the reward intrinsic value and predictability was manipulated. The bees could access a feeder whose sugar concentration varied from trial to trial following an increase, constant peak, and decrease pattern. Neither the feeder itself nor its position changed during the experiment. Hence, potential differences between trials could be interpreted in the context of decision-making and behavioural vigour, rather than search behaviour. The average flight speed towards the feeder increased when the sugar concentration was both high and constant, and returned to baseline as it became lower. This suggests that, as in other species including humans, the behavioural vigour of foraging bees towards a potential reward reflects its expected value. This motivational modulation of flight speed offer new ways of interpreting and studying decision-making in foraging bees.

## Introduction

Foraging animals assessing the value of a reward need to take into account both its inherent qualities (caloric, gustative, etc.) and the associated value modulators, such as its predictability and potential costs (1,2). A large body of research is dedicated to better understanding how each of these dimensions of value influences decision-making across animal species including humans (3–6).

An approach to measuring the value given by an experimental subject to a particular option is to compare sampling latencies (i.e. the time before the subject attends to an option when presented alone). Evidence that sampling latencies inversely correlates with the level of preference has accumulated in both avian and mammalian species (7,8). This correlation has its origin in the relationship between behavioural vigour and the average reward rate of the option available (9,10). In this context, behavioural vigour reflects the speed at which a decision is taken (11), the speed at which a motor behaviour is carried out (12,13); or a combination of both (10).

In bees, considerable research has been carried out on reward valuation during foraging (14– 18). As a result, how rewards affect flight behaviour has been studied from a variety of approaches, including optimal foraging theory (19,20) and the decision rules in foraging (21– 26), as well as the visual control of flight (27,28) and its energetic and performance aspects (29,30). Yet, while some of this research examined the time dimensions of decision-making (e.g. (31)), how reward affects behavioural vigour has received comparatively little attention. Among the few studies examining behavioural vigour in the context of reward valuation, honeybees have been shown to extend their proboscis faster after experiencing a series of increasingly large sugar rewards compared to constant or decreasing rewards (32), suggesting that the speed of decision-making is affected by variations in reward value. This study focused on the onset of behaviour, however, rather than its speed. Regarding the latter, the flight speed of forager bees leaving the hive has been shown to correlate with the concentration of sugar at the food source (33). More generally, an increase in reward flow rate generally leads to an increase in the frequency of visits to potential rewards (34–36). While these studies suggest a simple correlation between reward value and behavioural vigour, the relative importance of the reward intrinsic value and its value modulators is unclear. Moreover, two studies using judgment bias tasks have revealed that although an unexpected reward before a foraging flight reduces the sampling latencies of bees towards ambiguous stimuli, latency before choice to known options is unaffected (37,38). This suggests that changes in a bee’s motivational state do not systematically translate into changes in behavioural vigour.

The experiment presented here aims at further exploring this issue by examining whether a reward intrinsic value (sugar concentration) and predictability (similar concentration between consecutive trials) influence flight behaviour in free-flying common eastern bumblebees (*Bombus impatiens*). The flight parameters of foraging bees were recorded during ten foraging trials in which two consecutive high value trials were offered after an initial progressive increase in sugar concentration, and before a progressive decrease. A key aspect of this experimental paradigm is that the same feeder was used throughout the experiment, and its position was fixed. The feeder was also always rewarding in order to reduce the effects of search and acceptance factors on flight behaviour. This contrasts with other studies investigating the temporal aspects of decision-making in which latency before choice was influenced by the detectability and properties of the stimuli, or what bees had learned about them (e.g. (24,39–42)). Reducing the complexity of this classical foraging task was aimed to ensure that potential changes in flying behaviour could be interpreted as reflecting the effects of reward value and predictability on decision-making.

## Methods

### Subjects and arena

The nest box of a commercial bumblebee colony (Biobest North America, Leamington, ON, Canada) was connected to a flight arena made of white cardboard sheets supported by a wooden frame, and covered by a plastic mesh ceiling (figure 1). A separation wall with an opening at its top-centre (8 cm radius semi-circle, height: 50cm) divided the arena into the test arena and an intermediate arena. The role of this arena was to homogenise the flight parameters of the bees entering the test arena.

**Figure 1.**
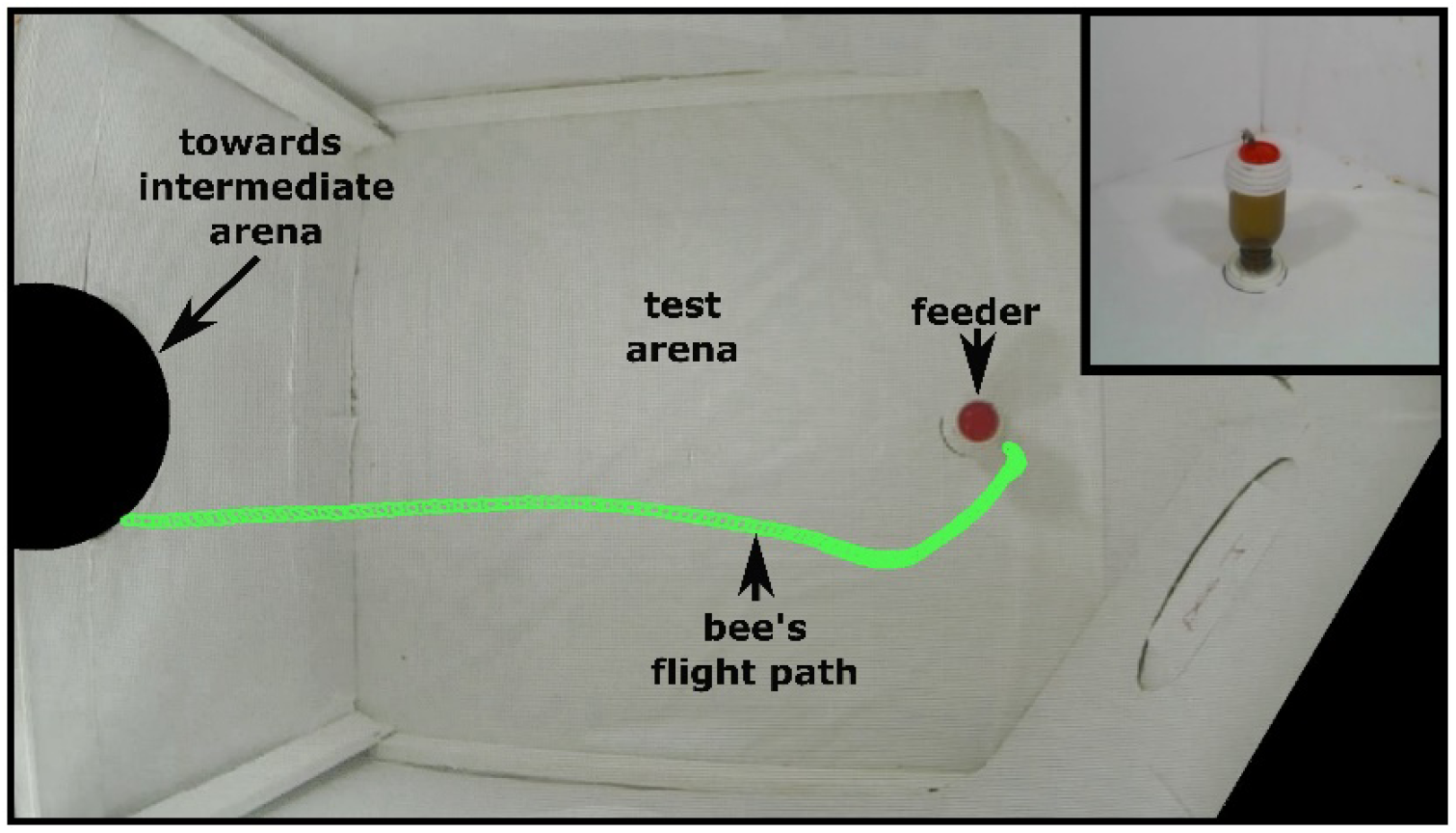
Annotated frame from the top camera used for the detection and tracking analysis. The flight path (in green) corresponds to the consecutive positions of the bee computed frame by frame. Masks (in black) were applied prior to video analysis to ensure that the bees were detected only when entering the test arena. Top right insert: view from the intermediate wall showing a bee feeding on the sugar solution.

### Feeder

The feeder was placed 56 cm away from the entrance of the test arena (figure 1). It was constituted of a 10 cm high upside-down brown tinted glass bottle with five loops of a white soft cotton rope (diameter 5 mm) glued around the top to ensure that the bees could easily grasp hold of the feeder when landing, from any direction of approach (figure 1). A red plastic dish 3 cm in diameter was placed at the top with around 0.5 ml of the appropriate sugar solution. The dish was replaced regularly during training to control for changes in sugar concentration through evaporation, and between each trials.

### Rewards

Five concentrations of sugar solution (in °Bx) were used in the following order: 30, 35, 40, 45, 50, 50, 45, 40, 35, 30. Bumblebees generally prefer the highest concentration within that range (43,44), such that sugar concentration can be considered here as reflecting the reward intrinsic value. As reward temperature influence preference in bumblebees (45), care was taken so that temperature was similar between the five sugar solutions (room temperature, ∼20°C).

### Predictability

The pattern of sugar concentration consisted in three consecutive phases: an increase, a constant peak, and a decrease. Since bees are sensitive to patterns of changes in reward quality (46), these phases cannot be considered as being totally unpredictable. However, because the response recorded in each trial depends on the value given by the bees to the feeder during the preceding trial(s), predictability differs between trials. The expected value of a trial may thus be: constant [trials 1 and 2] (due to the sugar concentration at trial 1 being similar to that during training), variable [trials 3 to 6] (due to the sugar concentration varying from trial to trial), constant [trial 7] (due to the two similar trials preceding it) and variable [trials 8 to 10] (due to the sugar concentration varying again from trial to trial). The pattern of rewards when both reward concentration and predictability is considered was thus: constant low, variable increasing, constant high, and variable decreasing.

### Videography

Trials were recorded with a GoPro Hero3 Black Edition (GoPro, Inc., 720*1280 px; 120 fps), and the bees’ position was computed frame by frame using a custom-made python program using OpenCV’s MOG2 (Mixture of Gaussian v2) algorithm (47) for background subtraction. Because the flight height was not measured, the position data is in pixel. Although pixels cannot be directly converted into a physical measure without additional data, the height of the camera relative to the floor of the arena remained unchanged during the experiment, which allows comparison between bees and conditions.

### Procedure

#### Training

Bees were first trained in group. Individual bees were gently captured with a small glass beaker before being brought into contact with the feeder containing a 30 °Bx sugar solution. Once several bees had started to forage autonomously, the separation wall was progressively raised until the bees could enter and exit the test arena through its opening. At this stage, the trained bees were paint marked so they could be individually identified and selected for the experiment.

#### Experiment

Twenty bees were given the ten trials described above in a continuous setting, one bee at a time. Each trial consisted in the tested bee entering the arena, foraging on the feeder, and going back to the nest. The experiment lasted around one hour per bee.

### Variables and analyses

Latency was obtained from the video files by subtracting the time at which a bee entered the arena from the time at which it landed on the feeder. Distance corresponds to the sum of the distance between consecutive positions. Mean speed was computed by averaging the individual speed data between consecutive positions. Maximum speed was obtained after fitting generalized additive models (GAM (48)) to the individual speed data in order to reduce the noise effect of localisation. Data from nine trials were lacking due to technical issues (n=3), bee falling into the feeder (n=1), or refusing to land on the feeder or enter the arena in trials 9 (n=2) and 10 (n=3).

### Statistical analysis

Linear mixed models (LMM) with trial as fixed factor and bee identity as random intercept were computed to analyse latency, average speed; distance and maximum speed. Thereafter, pairwise comparisons between trials were conducted for each variable for confirmation. The analyses were carried out both on the raw data and after removing the outliers with the following procedure: each variable was scanned and, for each bee, trials containing an absolute z-values greater than 3 were removed (seven trials from four bees). The main results are robust to the presence or absence of outliers, and the analyses reported here are based on the filtered data. All analyses were ran using R v4.2.1 (49).

### Results

#### Latency

The LMM analysis revealed a main effect of trial for latency (F(9,155.31)=3.38, p<0.001). More specifically, latency was significantly shorter in trial 7 compared to trials 1 to 6 as well as trial 10 (<0.001<p-values<0.048, 0.71>ds>1.47). Additionally, latency was shorter in trial 8 compared to latency at trials 2 to 4 and 6 (<0.001<p-values<0.038, 0.68>ds>1.15) as well as in trial 9 compared to trials 2 (p-value=0.004, d=0.97) and 4 (p-value=0.040, d=0.69). Finally, latency was longer in trial 2 compared to trials 5 (p-value=0.021, d=0.77) and 10 (p-value=0.024, d=0.78) (figure 2).

**Figure 2.**
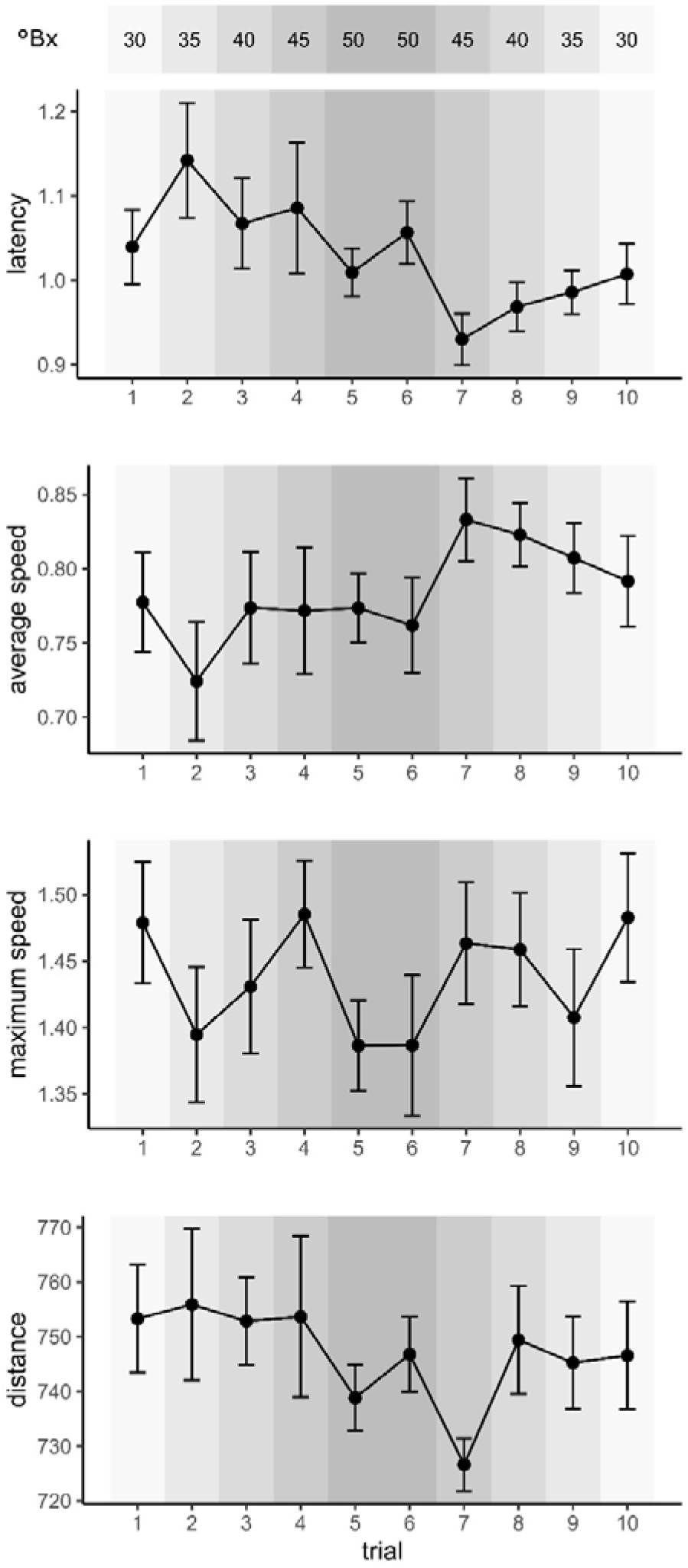
Flight parameters during the ten foraging trials (mean ± SE). The grey shading scale represents the concentration of sugar (in °Bx) for each trial. Latency is in seconds, distance in pixels and average/maximum speed in pixels/ms.

#### Average speed

The LMM analysis revealed a main effect of trial for average speed (F(9,155.17)=2.81, p=0.004). More specifically, average speed was significantly higher between trial 7 and trials 1 to 6 as well as in trial 10 (<0.001<p-values<0.033, 0.75>ds>1.36). Additionally, average speed of flight was higher in trial 8 compared to trials 2 to 6 (<0. 001<p-values<0.045, 0.66>ds>1.13), and in trial 9 compared to trial 2 (p-value=0.015, d=0.82) (figure 2).

#### Maximum speed

The LMM analysis revealed no significant effect of trial for the maximum speed within the arena (F(9,155.76)=1.09,p=0.374) (figure 2), which was corroborated by the follow up contrast analysis.

#### Distance

The LMM analysis revealed no significant effect of trial for distance travelled before landing (F(9,155.27)=1.50, p=0.153). However, follow-up contrasts revealed a shorter distance in trial 7 compared to trials 1 to 6 and 8 (0.003<p-values<0.025, 0.74>ds>1.02) (figure 2).

### Generalized Additive Models (GAM)

Visualization of the GAM estimates of the speed over the position of the bees on the axis entrance/feeder for each trial (figure 3), suggests that the difference in average speed in trial 7 compared to the other trials results from a lesser deceleration in the middle of the arena when approaching the feeder.

**Figure 3.**
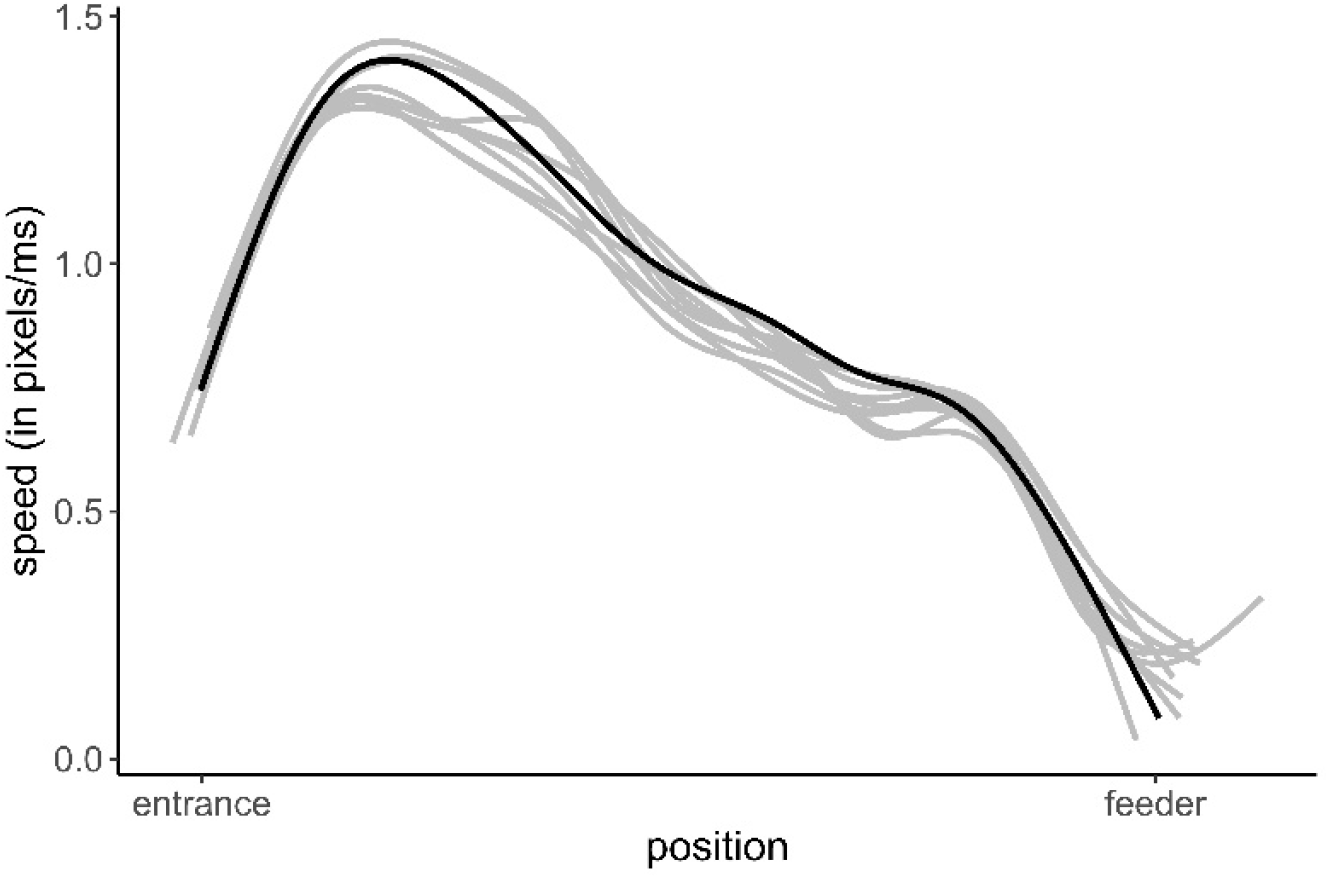
Average speed over the position on the axis entrance/feeder. Each line represents the GAM predictions on the data from all bees within a trial. Trial 7 is represented in black. The variability at the entrance and feeder positions is due to the semi-circular form of the entrance and the fact that bees could access the feeder from all sides.

## Discussion

The current study aimed at examining the effects of reward intrinsic value (sugar concentration) and predictability (similar concentration between consecutive trials) on flight behaviour in free-flying common eastern bumblebees (*Bombus impatiens*). The most significant finding of this experiment is the increase in average flight speed, and corresponding decrease in latency, at trial 7. This effect is unlikely to be an energetic (50,51) or social (52,53) consequence of the high concentration of sugar in the preceding trials, as average flight speed already diminishes at trial 8 despite the still relatively high concentration (45 °Bx) experienced at trial 7. In addition, an effect specific to the 50 °Bx reward solution would be expected to appear on trial 6, the second trial of the constant peak phase. Rather, this result suggests that flight speed increased when the reward value was both high and predictable (the bees could expect trial 7 to be the same as trials 5 and 6, both at 50 °Bx). Thus, like in other animals including humans (54), the expected value of a potential reward appears to influence the bees’ behavioural vigour (11,55–57).

This result complements an earlier observation by von Frisch and Lindauer (33) who reported a faster speed of departure in honey bees foraging on highly concentrated sugar. The novel finding here is that a potentially high intrinsic value alone is insufficient to induce changes in flight speed. Theoretical (9,10) and empirical (e.g. (58)) evidence suggests that behavioural vigour increases with average reward rate. Here, neither the increase in sugar concentration nor the two consecutive trials with the highest concentration led to a change in behavioural vigour. This is despite the fact that bees have been shown to form expectations about potential rewards (59,60), and alter their search behaviour based on their pattern of variation (46,61,62). This suggests that a reduction of uncertainty is required before the expectation of a reward alters flight speed in bees. Such an effect of reward predictability on the modulation of the wanting component of motivation seen here mirrors what has been reported in other animals including humans (63,64). This resemblance is rooted in the similarities between the neural reward systems of vertebrates and insects (65–68).

As shown in figure 3, the difference in average flight speed between trial 7 and the others seems to occur in the middle of the arena. Actually, both flying through the relatively narrow opening between the intermediate and test arenas and landing on the feeder involve stereotypical flight behaviours (69–71) that may limit the influence of motivational factors during these flight manoeuvres. In all cases, the generally higher flight speed in the middle of the arena suggests that the reduction in latency is due to a faster speed towards the feeder, rather than a faster reaction time. Indeed, foraging decisions seem to be partially taken before departure. This is suggested by the flexible intake of fuel load, which depends both on the predictability of the food source and the experience of the forager (72,73), and the variability, mentioned above, in the speed of departure from the hive, which depends on the quality of the expected reward (33).

The increase in behavioural vigour associated with the expectation of a high value reward can be understood as a way to avoid the cost of time in reward acquisition (6,10,74). In short, when the value of a reward is predicted to be high, any delay before its acquisition involves a cost that can be reduced by increasing behavioural vigour. In fact, the higher flight speed toward more valued potential rewards found here, and the resulting sampling bias due to lower search time (see also (41)), may be one of the factors responsible for the phenomenon of flower constancy found in several species of social bees (75). It could perhaps also help explain the “streaker bee” phenomenon (76,77), in which scout honeybees guide swarms by, among other things, flying faster in the direction of the potential nest site (78,79).

The progressive return to the initial average flying speed from trial 8 to 10 (during which sugar concentration decreased) contrasts with the abrupt increase in average flight speed in trial 7. In addition to corroborating the significance of the effect observed in trial 7 (by making it unlikely to result from simple effects of trial repetition), this pattern suggests that the inertia for change in reward valuation is asymmetric with regards to the direction of change (46,80). This asymmetry is partly caused by the non-independence of consecutive trials due to incentive contrasts (60,81), such that previously highly valued options become less preferred after the bees experience higher quality rewards. Importantly, the experiment does not allow to determine whether bees would maintain the elevated speed seen at trial 7 after several presentations of the highly valued reward, or return to the initial speed due to the costs associated with faster movements (30,82) (but see (83) for evidence of long lasting effects of high-value rewards on vigour).

Remarkably, trial 2 stands out compared to the relative constancy of the first six trials, particularly in terms of average speed. This is despite the fact that bees could have expected the sugar concentration to be the same as during training (30 °Bx). This suggests that unaccounted factors influenced the feeder valuation during the first trial (perhaps the absence of conspecifics or the longer wait in the selection tube before the experiment started). This also suggests that such a small-scale experimental set-up is sensitive enough to detect minor effects on flight speed, despite the high unrestricted flight speed of foraging bees (84).

The present results reveal a lack of effect of a liking component of reward processing on flight speed (see Ref. (85) for a review of the “liking” and “wanting” concepts of motivation). Such hedonic component is difficult to evaluate in bees, which lack the type of orofacial reactions commonly used in mammalian studies (66). While the study of antennal movements may offer a way to investigate such possible markers of liking (86–88), there is currently no accepted marker of “liking” in bees. The presence of a hedonic component is however suggested by the effect of unexpected rewards given to forager bees on their evaluation of ambiguous rewards (37,89). Here, such a “liking” effect as concentration increased did not produce significant changes in flight behaviour. While this is similar to what has been reported in experiments on cognitive bias tasks (37,38), the positive cognitive bias reported by these studies when bees are in a (supposedly) high hedonic state suggests that such a “liking” effect on flight behaviour is possible, perhaps in terms of trajectory (which betrays the search of alternatives within the environment). Importantly, by virtue of happening before reward consumption, any behavioural change in the flight characteristics of a bee flying toward a reward could be interpreted not as evidence of liking, but rather of the “expected pleasantness” of the reward (90).

Experimental protocols used to study reward valuation and decision-making in foraging bees classically consist in giving (classically, tethered (91)) bees a choice between rewarding and unrewarding, or aversive, options (92,93), or between options varying in the quality of the reward offered or their probability of being rewarding (61,94–96). Two often overlooked factors are the time dimension of decision-making (97) and, as shown here, behavioural vigour. The present results suggest that studying the flight parameters of foraging bees, which involves both of these factors, may help study the different valuation subsystems involved in decision-making (see also (40)). By dissociating the effects of the liking and wanting components of motivation, this type of analysis offers a potential way forward to a long lasting problem in insect cognitive studies (66). Standard recordings of behavioural vigour via high-speed videography (see also (98)) could thus provide an alternative, or complementary way of measuring preference between options, beyond the multi-choice trials classically used (8). Knowledge could be gained not only by examining the factors leading to a change in the flight parameters, but also by studying the rate of decline of these parameters as the expectations regarding a simple or composed (99) potential reward are altered.

## Notes

### Competing Interest Statement

The authors have declared no competing interest.

## References

1. Basten U, Biele G, Heekeren HR, Fiebach CJ. How the brain integrates costs and benefits during decision making. Proceedings of the National Academy of Sciences. National Acad Sciences; 2010;107(50):21767–21772.

2. Rangel A, Camerer C, Montague PR. A framework for studying the neurobiology of value-based decision making. Nature reviews neuroscience. Nature Publishing Group UK London; 2008;9(7):545–556.

3. Dall SR, Giraldeau L-A, Olsson O, McNamara JM, Stephens DW. Information and its use by animals in evolutionary ecology. Trends in ecology & evolution. Elsevier; 2005;20(4):187–193.

4. Doya K. Modulators of decision making. Nature neuroscience. Nature Publishing Group US New York; 2008;11(4):410–416.

5. Schultz W. Multiple reward signals in the brain. Nature reviews neuroscience. Nature Publishing Group UK London; 2000;1(3):199–207.

6. Yoon T, Geary RB, Ahmed AA, Shadmehr R. Control of movement vigor and decision making during foraging. Proceedings of the National Academy of Sciences. National Acad Sciences; 2018;115(44):E10476–E10485.

7. Elliott MH. The effect of change of reward on the maze performance of rats. University of California Press; 1928.

8. Kacelnik A, Vasconcelos M, Monteiro T. Testing cognitive models of decision-making: selected studies with starlings. Animal Cognition. Springer; 2023;26(1):117–127.

9. Cools R, Nakamura K, Daw ND. Serotonin and dopamine: unifying affective, activational, and decision functions. Neuropsychopharmacology. Nature Publishing Group; 2011;36(1):98–113.

10. Niv Y, Daw ND, Joel D, Dayan P. Tonic dopamine: opportunity costs and the control of response vigor. Psychopharmacology. Springer; 2007;191:507–520.

11. Shapiro MS, Siller S, Kacelnik A. Simultaneous and sequential choice as a function of reward delay and magnitude: normative, descriptive and process-based models tested in the European starling (Sturnus vulgaris). Journal of Experimental Psychology: Animal Behavior Processes. American Psychological Association; 2008;34(1):75.

12. Crespi LP. Quantitative variation of incentive and performance in the white rat. The American Journal of Psychology. JSTOR; 1942;55(4):467–517.

13. Shadmehr R, De Xivry JJO, Xu-Wilson M, Shih T-Y. Temporal discounting of reward and the cost of time in motor control. Journal of Neuroscience. Soc Neuroscience; 2010;30(31):10507–10516.

14. Bitterman ME. Comparative analysis of learning in honeybees. Animal Learning & Behavior. Springer; 1996;24(2):123–141.

15. McNeill M, Kapheim K, Brockmann A, McGill T, Robinson G. Brain regions and molecular pathways responding to food reward type and value in honey bees. Genes, Brain and Behavior. Wiley Online Library; 2016;15(3):305–317.

16. Núñez JA, Giurfa M. Motivation and regulation of honey bee foraging. Bee World. Taylor & Francis; 1996;77(4):182–196.

17. Pleasants JM. Bumblebee response to variation in nectar availability. Ecology. Wiley Online Library; 1981;62(6):1648–1661.

18. Seeley TD. Social foraging by honeybees: how colonies allocate foragers among patches of flowers. Behavioral Ecology and Sociobiology. Springer; 1986;19:343–354.

19. Núñez J. Honeybee foraging strategies at a food source in relation to its distance from the hive and the rate of sugar flow. Journal of Apicultural Research. Taylor & Francis; 1982;21(3):139–150.

20. Schmid-Hempel P, Kacelnik A, Houston AI. Honeybees maximize efficiency by not filling their crop. Behavioral Ecology and Sociobiology. Springer; 1985;17(1):61–66.

21. Chittka L, Gumbert A, Kunze J. Foraging dynamics of bumble bees: correlates of movements within and between plant species. Behavioral Ecology. Oxford University Press; 1997;8(3):239–249.

22. Greggers U, Menzel R. Memory dynamics and foraging strategies of honeybees. Behavioral Ecology and Sociobiology. Springer; 1993;32:17–29.

23. Kadmon R, Shmida A. Departure rules used by bees foraging for nectar: a field test. Evolutionary Ecology. Springer; 1992;6:142–151.

24. Keasar T, Shmida A, Motro U. Innate movement rules in foraging bees: flight distances are affected by recent rewards and are correlated with choice of flower type. Behavioral Ecology and Sociobiology. Springer; 1996;39:381–388.

25. Lihoreau M, Chittka L, Raine NE. Trade-off between travel distance and prioritization of high-reward sites in traplining bumblebees. Functional ecology. Wiley Online Library; 2011;25(6):1284–1292.

26. Schmid-Hempel P. The influence of reward sequence on flight directionality in bees. Animal behaviour. Elsevier; 1986;34(3):831–837.

27. Baird E, Srinivasan MV, Zhang S, Cowling A. Visual control of flight speed in honeybees. Journal of experimental biology. The Company of Biologists Ltd; 2005;208(20):3895–3905.

28. Si A, Srinivasan MV, Zhang S. Honeybee navigation: properties of the visually drivenodometer. Journal of Experimental Biology. The Company of Biologists Ltd; 2003;206(8):1265–1273.

29. Burnett NP, Badger MA, Combes SA. Wind and obstacle motion affect honeybee flight strategies in cluttered environments. Journal of Experimental Biology. The Company of Biologists Ltd; 2020;223(14):jeb222471.

30. Nachtigall W, Hanauer-Thieser U, Mörz M. Flight of the honey bee VII: metabolic power versus flight speed relation. Journal of Comparative Physiology B. Springer; 1995;165(6):484–489.

31. Skorupski P, Spaethe J, Chittka L. Visual search and decision making in bees: time, speed, and accuracy. International Journal of Comparative Psychology. 2006;19(3).

32. Gil M, Menzel R, De Marco RJ. Does an insect’s unconditioned response to sucrose reveal expectations of reward? PLoS One. Public Library of Science San Francisco, USA; 2008;3(7):e2810.

33. Frisch K v, Lindauer M. Über die Fluggeschwindigkeit der Bienen und über ihre Richtungsweisung bei Seitenwind. Naturwissenschaften. Springer; 1955;42:377–385.

34. Balderrama N, Almeida L de, Núñez J. Metabolic rate during foraging in the honeybee. Journal of Comparative Physiology B. Springer; 1992;162:440–447.

35. Giurfa M. Movement patterns of honeybee foragers: motivation and decision rules dependent on the rate of reward. Behaviour. JSTOR; 1996;579–596.

36. Giurfa M, Núñez J a. Foraging by honeybees on Carduus acanthoides: pattern and efficiency. Ecological Entomology. Wiley Online Library; 1992;17(4):326–330.

37. Perry CJ, Baciadonna L, Chittka L. Unexpected rewards induce dopamine-dependent positive emotion–like state changes in bumblebees. Science. American Association for the Advancement of Science; 2016;353(6307):1529–1531.

38. Strang C, Muth F. Judgement bias may be explained by shifts in stimulus response curves. Royal Society Open Science. The Royal Society; 2023;10(4):221322.

39. Chittka L, Dyer AG, Bock F, Dornhaus A. Psychophysics: bees trade off foraging speed for accuracy. Nature. Nature Publishing Group; 2003;424(6947):388.

40. MaBouDi H, Marshall JA, Dearden N, Barron AB. How honey bees make fast and accurate decisions. Grunwald Kadow IC, Frank MJ, Riabinina O, Fujiwara T, editors. eLife [Internet]. eLife Sciences Publications, Ltd; 2023 Jun;12:e86176. Available from: 10.7554/eLife.86176

41. Nityananda V, Chittka L. Different effects of reward value and saliency during bumblebee visual search for multiple rewarding targets. Animal Cognition. Springer; 2021;24(4):803–814.

42. Spaethe J, Tautz J, Chittka L. Visual constraints in foraging bumblebees: flower size and color affect search time and flight behavior. Proceedings of the National Academy of Sciences. National Acad Sciences; 2001;98(7):3898–3903.

43. Bailes EJ, Pattrick JG, Glover BJ. An analysis of the energetic reward offered by field bean (Vicia faba) flowers: Nectar, pollen, and operative force. Ecology and evolution. Wiley Online Library; 2018;8(6):3161–3171.

44. Konzmann S, Lunau K. Divergent rules for pollen and nectar foraging bumblebees–a laboratory study with artificial flowers offering diluted nectar substitute and pollen surrogate. PLoS one. Public Library of Science San Francisco, USA; 2014;9(3):e91900.

45. Dyer AG, Whitney HM, Arnold SE, Glover BJ, Chittka L. Bees associate warmth with floral colour. Nature. Nature Publishing Group UK London; 2006;442(7102):525–525.

46. Gil M. Reward expectations in honeybees. Communicative & Integrative Biology. Taylor & Francis; 2010;3(2):95–100.

47. Zivkovic Z, Van Der Heijden F. Efficient adaptive density estimation per image pixel for the task of background subtraction. Pattern recognition letters. Elsevier; 2006;27(7):773–780.

48. Wood S, Wood MS. Package ‘mgcv. R package version. 2015;1(29):729.

49. Team RC. R: A Language and Environment for Statistical Computing [Internet]. Vienna, Austria: R Foundation for Statistical Computing; 2022. Available from: https://www.R-project.org/

50. Nieh JC, León A, Cameron S, Vandame R. Hot bumble bees at good food: thoracic temperature of feeding Bombus wilmattae foragers is tuned to sugar concentration. Journal of Experimental Biology. Company of Biologists; 2006;209(21):4185–4192.

51. Stabentheiner A, Hagmüller K. Sweet food means” hot dancing” in honeybees. Naturwissenschaften. 1991;78:471–473.

52. Cartar RV. Adjustment of foraging effort and task switching in energy-manipulated wild bumblebee colonies. Animal Behaviour. Elsevier; 1992;44(1):75–87.

53. Franklin EL, Smith KE, Raine NE. How foraging preference and activity level of bumble bees contribute to colony flexibility under resource demand. Animal Behaviour. Elsevier; 2022;194:43–55.

54. Montague PR, Dayan P, Sejnowski TJ. A framework for mesencephalic dopamine systems based on predictive Hebbian learning. Journal of neuroscience. Soc Neuroscience; 1996;16(5):1936–1947.

55. Monteiro T, Vasconcelos M, Kacelnik A. Choosing fast and simply: Construction of preferences by starlings through parallel option valuation. PLoS Biology. Public Library of Science San Francisco, CA USA; 2020;18(8):e3000841.

56. Roces F. Individual complexity and self-organization in foraging by leaf-cutting ants. The Biological Bulletin. Marine Biological Laboratory; 2002;202(3):306–313.

57. Shadmehr R, Reppert TR, Summerside EM, Yoon T, Ahmed AA. Movement vigor as a reflection of subjective economic utility. Trends in neurosciences. Elsevier; 2019;42(5):323–336.

58. Guitart-Masip M, Beierholm UR, Dolan R, Duzel E, Dayan P. Vigor in the face of fluctuating rates of reward: an experimental examination. Journal of cognitive neuroscience. MIT Press One Rogers Street; 2011;23(12):3933–3938.

59. Couvillon P, Bitterman M. The overlearning-extinction effect and successive negative contrast in honeybees (Apis mellifera). Journal of Comparative Psychology. American Psychological Association; 1984;98(1):100.

60. Hemingway CT, Muth F. Label-based expectations affect incentive contrast effects in bumblebees. Biology Letters. 2022;18(3).

61. Drezner-Levy T, Shafir S. Parameters of variable reward distributions that affect risk sensitivity of honey bees. Journal of Experimental Biology. Company of Biologists; 2007;210(2):269–277.

62. Gil M, De Marco RJ, Menzel R. Learning reward expectations in honeybees. Learning & Memory. Cold Spring Harbor Laboratory Press; 2007;14(7):491.

63. Berns GS, McClure SM, Pagnoni G, Montague PR. Predictability modulates human brain response to reward. Journal of neuroscience. Soc Neuroscience; 2001;21(8):2793–2798.

64. Webber HE, Lopez-Gamundi P, Stamatovich SN, Wit H de, Wardle MC. Using pharmacological manipulations to study the role of dopamine in human reward functioning: A review of studies in healthy adults. Neuroscience & Biobehavioral Reviews. Elsevier; 2021;120:123–158.

65. Huang J, Zhang Z, Feng W, Zhao Y, Aldanondo A, Brito Sanchez MG de, et al. Food wanting is mediated by transient activation of dopaminergic signaling in the honey bee brain. Science. American Association for the Advancement of Science; 2022;376(6592):508–512.

66. Perry CJ, Barron AB. Neural mechanisms of reward in insects. Annual review of entomology. Annual Reviews; 2013;58:543–562.

67. Scheiner R, Baumann A, Blenau W. Aminergic control and modulation of honeybee behaviour. Current neuropharmacology. Bentham Science Publishers; 2006;4(4):259– 276.

68. Wright G. The role of dopamine and serotonin in conditioned food aversion learning in the honeybee. Communicative & integrative biology. Taylor & Francis; 2011;4(3):318– 320.

69. Baird E, Boeddeker N, Ibbotson MR, Srinivasan MV. A universal strategy for visually guided landing. Proceedings of the National Academy of Sciences. National Acad Sciences; 2013;110(46):18686–18691.

70. Reber T, Baird E, Dacke M. The final moments of landing in bumblebees, Bombus terrestris. Journal of Comparative Physiology A. Springer; 2016;202:277–285.

71. Srinivasan M, Zhang S, Lehrer M, Collett T. Honeybee navigation en route to the goal: visual flight control and odometry. Journal of Experimental Biology. The Company of Biologists Ltd; 1996;199(1):237–244.

72. Harano K, Mitsuhata-Asai A, Konishi T, Suzuki T, Sasaki M. Honeybee foragers adjust crop contents before leaving the hive. Behavioral Ecology and Sociobiology. Springer; 2013;67(7):1169–1178.

73. Tan K, Latty T, Dong S, Liu X, Wang C, Oldroyd BP. Individual honey bee (Apis cerana) foragers adjust their fuel load to match variability in forage reward. Scientific reports. Nature Publishing Group; 2015;5:16418.

74. Berret B, Jean F. Why don’t we move slower? The value of time in the neural control of action. Journal of neuroscience. Soc Neuroscience; 2016;36(4):1056–1070.

75. Grüter C, Ratnieks FL. Flower constancy in insect pollinators. Communicative & Integrative Biology. 2011;4(6):633–636.

76. Beekman M, Fathke RL, Seeley TD. How does an informed minority of scouts guide a honeybee swarm as it flies to its new home? Animal behaviour. Elsevier; 2006;71(1):161–171.

77. Lindauer M. Schwarmbienen auf wohnungssuche. Zeitschrift für vergleichende Physiologie. Springer; 1955;37:263–324.

78. Greggers U, Schoening C, Degen J, Menzel R. Scouts behave as streakers in honeybee swarms. Naturwissenschaften. Springer; 2013;100:805–809.

79. Schultz KM, Passino KM, Seeley TD. The mechanism of flight guidance in honeybee swarms: subtle guides or streaker bees? Journal of Experimental Biology. Company of Biologists; 2008;211(20):3287–3295.

80. Wainselboim AJ, Roces F, Farina WM. Honeybees assess changes in nectar flow within a single foraging bout. Animal Behaviour. Elsevier; 2002;63(1):1–6.

81. Bitterman M. Incentive contrast in honey bees. Science. American Association for the Advancement of Science; 1976;192(4237):380–382.

82. Berret B, Baud-Bovy G. Evidence for a cost of time in the invigoration of isometric reaching movements. Journal of Neurophysiology. American Physiological Society Rockville, MD; 2022;127(3):689–701.

83. Wagner AR. Effects of amount and percentage of reinforcement and number of acquisition trials on conditioning and extinction. Journal of experimental Psychology. American Psychological Association; 1961;62(3):234.

84. Baracchi D, Lihoreau M, Giurfa M. Do insects have emotions? Some insights from bumble bees. Frontiers in Behavioral Neuroscience. Frontiers Media SA; 2017;11:157.

85. Berridge KC, Robinson TE, Aldridge JW. Dissecting components of reward:”liking”,”wanting”, and learning. Current opinion in pharmacology. Elsevier; 2009;9(1):65–73.

86. Cholé H, Merlin A, Henderson N, Paupy E, Mahé P, Arnold G, et al. Antenna movements as a function of odorants’ biological value in honeybees (Apis mellifera L.). Scientific Reports. Nature Publishing Group UK London; 2022;12(1):11674.

87. Claverie N, Buvat P, Casas J. Active sensing in bees through antennal movements is independent of odor molecule. Integrative and Comparative Biology. Oxford University Press; 2023;63(2):315–331.

88. Gascue F, Marachlian E, Azcueta M, Locatelli FF, Klappenbach M. Antennal movements can be used as behavioral readout of odor valence in honey bees. IBRO Neuroscience Reports. Elsevier; 2022;12:323–332.

89. Perry CJ, Barron AB, Chittka L. The frontiers of insect cognition. Current Opinion in Behavioral Sciences. Elsevier; 2017;16:111–118.

90. Pool E, Sennwald V, Delplanque S, Brosch T, Sander D. Measuring wanting and liking from animals to humans: A systematic review. Neuroscience & Biobehavioral Reviews. Elsevier; 2016;63:124–142.

91. Giurfa M, Sandoz J-C. Invertebrate learning and memory: fifty years of olfactory conditioning of the proboscis extension response in honeybees. Learning & memory. Cold Spring Harbor Lab; 2012;19(2):54–66.

92. Avarguès-Weber A, Brito Sanchez MG de, Giurfa M, Dyer AG. Aversive reinforcement improves visual discrimination learning in free-flying honeybees. PLoS One. Public Library of Science; 2010;5(10):e15370.

93. Muth F, Cooper TR, Bonilla RF, Leonard AS. A novel protocol for studying bee cognition in the wild. Methods in Ecology and Evolution. Wiley Online Library; 2018;9(1):78–87.

94. Anselme P. Uncertainty processing in bees exposed to free choices: Lessons from vertebrates. Psychonomic Bulletin & Review. Springer; 2018;25:2024–2036.

95. Benard J, Giurfa M. A test of transitive inferences in free-flying honeybees: unsuccessful performance due to memory constraints. Learning & Memory. Cold Spring Harbor Lab; 2004;11(3):328–336.

96. MaBouDi H, Marshall JA, Barron AB. Honeybees solve a multi-comparison ranking task by probability matching. Proceedings of the Royal Society B. The Royal Society; 2020;287(1934):20201525.

97. Chittka L, Spaethe J. Visual search and the importance of time in complex decision making by bees. Arthropod-Plant Interactions. Springer; 2007;1:37–44.

98. Guiraud M, Roper M, Chittka L. High speed videography reveals how honeybees can turn a spatial concept learning task into a simple discrimination task by stereotyped flight movements and sequential inspection of pattern elements. Frontiers in psychology. Frontiers; 2018;9:1347.

99. Muth F, Papaj DR, Leonard AS. Multiple rewards have asymmetric effects on learning in bumblebees. Animal Behaviour. Elsevier; 2017;126:123–133.

